# Advances in Estimating Level-1 Phylogenetic Networks from Unrooted SNPs

**DOI:** 10.1101/2024.07.19.604386

**Authors:** Tandy Warnow, Yasamin Tabatabaee, Steven N. Evans

## Abstract

We address the problem of how to estimate a phylogenetic network when given SNPs (i.e., single nucleotide polymorphisms, or bi-allelic markers that have evolved under the infinite sites assumption). We focus on level-1 phylogenetic networks (i.e., networks where the cycles are node-disjoint), since more complex networks are unidentifiable. We provide a polynomial time quartet-based method that we prove correct for reconstructing the unrooted topology of any level-1 phylogenetic network *N*, if we are given a set of SNPs that covers all the bipartitions of *N*, even if the ancestral state is not known, provided that the cycles are of length at least 5; we also prove that an algorithm developed by Dan Gusfield in JCSS 2005 correctly recovers the unrooted topology in polynomial time in this case. To the best of our knowledge, this is the first result to establish that the unrooted topology of a level-1 network is uniquely recoverable from SNPs without known ancestral states. We also present a stochastic model for DNA evolution, and we prove that the two methods (our quartet-based method and Gusfield’s method) are statistically consistent estimators of the unrooted topology of the level-1 phylogenetic network. For the case of multi-state homoplasy-free characters, we prove that our quartet-based method correctly constructs the unrooted topology of level-1 networks under the required conditions (all cycles of length at least five), while Gusfield’s algorithm cannot be used in that condition. These results assume that we have access to an oracle for indicating which sites in the DNA alignment are homoplasy-free, and we show that the methods are robust, under some conditions, to oracle errors.

## 1 Introduction

The inference of phylogenetic trees from multiple sequence alignments is a basic step in much biological research, including the detection of selection, dating of diversifications, understanding biomolecular function and structure, etc. Yet, phylogenetic trees are simplistic models of evolution that do not allow for more complex events, such as horizontal gene transfers (Soucy et al., 2015), recombination (Schierup and Hein, 2000), and hybridization (Grant and Grant, 1992). Given the frequency of such events, more complex graphical models of evolutionary histories are needed (Morrison, 2005; Hernandez-Lopez, 2013). These models, which have additional edges compared to phylogenetic trees, are referred to as “phylogenetic networks”, and several books have been written on the subject (Huson et al., 2010; Morrison, 2011; Gusfield, 2014).

One of the applications of phylogenetic networks is in the context of population genetics, where the assumption is that certain sites in a multiple sequence alignment evolve under the infinite sites assumption, and so will evolve without any homoplasy and exhibit only two states. These sites, referred to as “single nucleotide polymorphisms”, or SNPs, can then be used to estimate the evolutionary history for the population (Gusfield, 2005, 2014).

When the graphical model for evolution is a tree (i.e., there is no reticulation in the evolution), then the tree is referred to as a “perfect phylogeny” and the characters are said to be “compatible” (Felsenstein, 1982). The estimation of perfect phylogenies from SNPs is easily seen to be polynomial time, even when the ancestral state is unknown, and the minimally resolved perfect phylogeny is unique (Gusfield, 1991). However, when recombination or other reticulate events occur, then more complex models, such as level-1 phylogenetic networks (i.e., networks where all cycles are node-disjoint, see Figure 1), are needed to represent the evolutionary history. In this context, each SNP is assumed to evolve under the infinite sites assumption down some rooted tree contained in the rooted phylogenetic network, and the problem is then to infer the phylogenetic network from the SNPs. Methods for constructing these networks have been developed, both for the case where the ancestral state is known as well as for the case where the ancestral state is unknown (Gusfield, 2005). However, it is not known whether these methods are statistically consistent under any formal stochastic model of SNP evolution down a phylogenetic tree or network, which is the main focus of this study.

**Figure 1:**
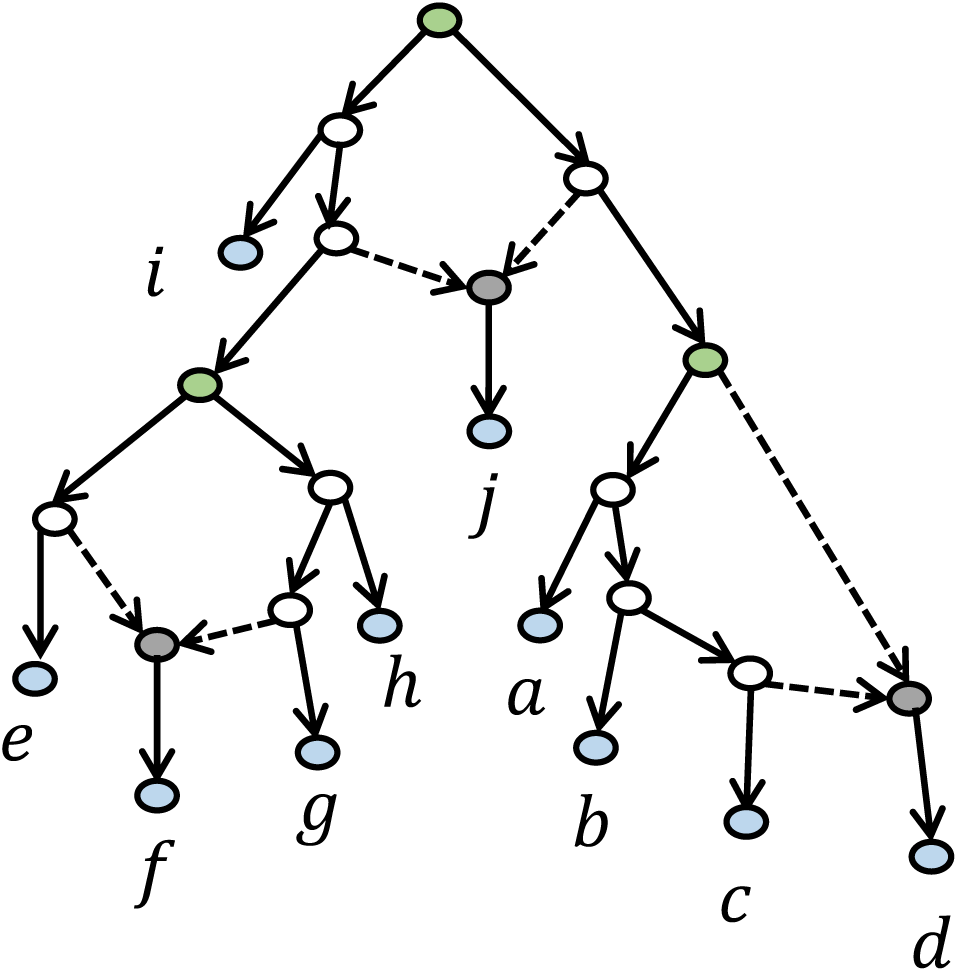
A phylogenetic network with three cycles. Since the three cycles do not share any nodes, this is a level-1 phylogenetic network. Each cycle has one node with indegree two, which makes it a reticulate node; these are indicated in gray, and the two incoming edges are dashed to indicate they are reticulate edges. Each cycle also has one node with two children within the cycle (the root of the cycle, indicated in green).

In practice, not all sites within a multiple sequence alignment will satisfy the requirements for SNPs: being two-state and having evolved under the infinite sites assumption (i.e., without homoplasy). Therefore, in practice, the biologist will use some tests to determine which sites are likely to be SNPs, and then use those sites in the estimation of the phylogenetic network.

In general, the estimation of phylogenetic networks is very difficult. One aspect that makes it challenging is that unlike phylogenetic trees, phylogenetic networks can have unbounded complexity, making for non-uniqueness of the underlying graphical model. Level-1 phylogenetic networks, defined above, represent a very simple graphical model that allows for reticulation but requires that no two cycles share any nodes in common. These rooted phylogenetic networks, referred to also as “galled trees”, are uniquely defined by their rooted triplet trees, their clades, or by the set of rooted trees contained within the rooted network, provided that no cycle in the network is of length less than five (Habib and To, 2012; Huber et al., 2010; Jansson et al., 2006; Poormohammadi et al., 2014; Van Iersel et al., 2009, 2010).

Of these algorithms, the fastest algorithm for constructing a level-1 network *N* from its set of triplet trees is the *O*(*n*^3^) algorithm given in Jansson et al. (2006), where *n* is the number of leaves in *N* . Hence, given the set *Clades*(*N*) of the clades of a network *N*, it is possible to construct the rooted topology of *N* by computing the set *Triplets*(*N*) from *Clades*(*N*) in *O*(*n*^4^) time, and then applying the algorithm from Jansson et al. (2006).

Since SNPs are assumed to evolve down trees contained within the level-1 phylogenetic network, it is also possible to construct networks from SNPs, as long as the ancestral state is known for each SNP. Specifically, if the SNP has states 0 and 1, where 0 is the ancestral state, then the leaves exhibiting state 1 form a clade in one of the trees within the network. Hence, these SNPs can be used to define clades, and then clade-based methods can be used to construct the network. A particularly fast method of this type is in Gusfield et al. (2003), which presents an *O*(*mn* + *n*^3^) algorithm to construct the level-1 network from *m* SNPs that cover the clades of the network, provided that the ancestral state is known, allowing for multiple crossover events, and assuming all cycles are of length at least five. It is easy to see that the number of clades in a level-1 network with *n* leaves is *O*(*n*), but see Gambette et al. (2017) (Section 4.1) for a concrete upper bound. Hence, *m* is *O*(*n*), and so the *O*(*mn* + *n*^3^) algorithm for constructing the level-1 network from *m* SNPs in turn implies an *O*(*n*^3^) algorithm to construct the network from its set of clades.

The construction of more complex phylogenetic networks may not be feasible: for example, slightly more complex networks, referred to as level-2 phylogenetic networks, can fail to be recovered from their triplets and clades, as demonstrated in Gambette and Huber (2012). For this reason, the research community in phylogenetic networks has largely restricted attention to constructing level-1 phylogenetic networks.

However, relatively little is understood about estimating level-1 phylogenetic networks from SNPs when the ancestral state is unknown (this is also referred to as the “root unknown” condition). That is, when the ancestral state is not provided, the SNPs do not define clades and instead define bipartitions, and hence methods that rely on having clades (or rooted triplet trees) are inapplicable. Furthermore, in practice, the ancestral state for each SNP is typically not known. Hence, understanding how to estimate phylogenetic networks from SNPs where the ancestral state is unknown is the biologically relevant problem.

Two other studies are relevant to the case where the SNPs are provided without information about the ancestral state Gusfield (2005), which assumes the cycles are node-disjoint, and Huson and Kloepper (2007), which allows cycles to share nodes but not edges. Here we focus on Gusfield’s contribution, as it is particularly computationally efficient (running in *O*(*mn* + *n*^3^) time, where there are *n* leaves and *m* bipartitions). Gusfield’s algorithm has the guarantee that if the input SNPS cover all the bipartitions of a level-1 phylogenetic network and each cycle has length at least five, then his algorithm will output an unrooted level-1 phylogenetic network that will be *consistent* with the input SNPs. Neverthleless, Gusfield did not establish that the output network would have the same unrooted topology as the true network, and left open the possibility that there could be two or more unrooted level-1 phylogenetic networks consistent with the input SNPs. .

Another contribution that is relevant is Gambette et al. (2012). In their study, they showed that the set *Q*(*N*) of all quartet trees of level-1 phylogenetic network *N* is sufficient to define the unrooted topology of *N*, where *Q*(*N*) contains all quartet trees *AB*|*CD* (for four leaves *A, B, C, D*) where there are node-disjoint paths *P_AB_* connecting *A* and *B* and *P_CD_* connecting *C* and *D* in *N* . Furthermore, they provided a polynomial time algorithm to construct the unrooted topology of *N* from *Q*(*N*). However, it is not clear that this approach can be used to construct unrooted level-1 phylogenetic networks from SNPs, since it is not known if the SNPs provide all the quartet trees in *Q*(*N*).

In this paper, we make several contributions for constructing level-1 phylogenetic networks from SNPs. First, we show that whenever the level-1 network contains any cycle of length at least five, the set *Q*(*N*) will contain quartet trees that are not derivable from SNPs, and the method from Gambette et al. (2012) will fail to reconstruct *N*, even when given SNPs that cover all the bipartitions of *N* . Thus, the method from Gambette et al. (2012) will fail to have desirable theoretical guarantees for constructing the unrooted level-1 phylogenetic network from SNPs.

The second major contribution is a new polynomial time method, CUPNS (Constructing Unrooted Phylogenetic Networks from SNPs), that is guaranteed to correctly construct the unrooted topology of any level-1 phylogenetic network *N* whose cycles are all of length at least five, when given SNPs without known ancestral state that cover the bipartitions of *N* . CUPNS is a quartet-based method that operates by computing quartet trees *AB*|*CD* where *A* and *B* share one state and *C* and *D* share the other state, and uses these quartet trees to construct *N* . This set of quartet trees can be seen as quartet trees found in rooted trees within the rooted phylogenetic network, and so is a subset *Q*(*N_r_*) of the set *Q*(*N*). As a consequence, we establish that *Q*(*N_r_*) is sufficient to define the unrooted level-1 phylogenetic network *N* . As a corollary, we establish that the algorithm in Gusfield (2005) provably correctly constructs the unrooted topology of the true level-1 phylogenetic network *N* when the SNPs cover the bipartitions of *N* and the cycles are of length at least five.

We then propose a stochastic model of site evolution within a level-1 phylogenetic network *N*, in which some sites are SNPs. We show that both CUPNS and Gusfield’s algorithm are statistically consistent estimators of the unrooted topology of *N* under this model (even without knowing the ancestral state for any SNP), provided that we have access to an oracle that correctly identifies the SNPs (i.e., two-state characters that are homoplasy-free), and the network cycles are all of length at least five.

The rest of the paper is organized as follows. We begin with basic material in Section 2. We then provide the new theory for computing the unrooted topology of a level-1 phylogenetic network *N* from *Q*(*N_r_*) in Section 3; this is the section where we present the CUPNS method. In Section 4 we present our stochastic model of DNA site evolution, and discuss statistical properties (e.g., statistical consistency) of different pipelines for estimating level-1 phylogenetic networks under this model, assuming availability of a perfect oracle. In Section 5, we discuss the limitations and extensions of the theory we have established, specifically examining robustness to assumptions. We conclude in Section 6 with a discussion of the ramifications of the theory and suggested directions for future work.

## 2 Preliminary material

### 2.1 Phylogenetic trees and networks

We begin with a description of the basic graphical models of evolution: phylogenetic trees and phylogenetic networks, as well as terminology associated to each.

#### Phylogenetic trees

Phylogenetic trees are rooted binary trees with leaves labelling the given set of taxa. The edges are directed away from the root, so that the unique root has indegree 0 and outdegree 2, the leaves have indegree 1 and outdegree 0, and all other nodes have indegree 1 and outdegree 2. The phylogenetic tree model is the standard graphical model of evolution, but does not allow for more complex events such as hybridization, lateral transfer, or recombination.

#### Phylogenetic networks

To address this limitation, phylogenetic network models were developed. Since a phylogenetic network is a graphical model of evolutionary history, there is a unique root, and all the edges are oriented away from the root, making the network a rooted directed acylic graph (DAG).

The unique root has indegree 0 and the leaves have indegree 1 and outdegree 0. The other nodes are either “tree nodes” or “reticulate nodes”. The tree nodes have indegree 1 and so resemble nodes in trees, while the reticulate nodes have indegree 2. These reticulate nodes may reflect hybridization, targets of lateral transfer, or other reticulation events. As with a phylogenetic tree, a fully resolved phylogenetic network would have outdegree 2 for the tree nodes and outdegree 1 for the reticulate nodes; however, for this study we allow tree nodes to have outdegrees at least two and reticulate nodes to have any positive outdegree.

Also, to further distinguish a phylogenetic network from a phylogenetic tree, the edges of the network are denoted either as tree edges or as reticulate edges. Specifically, any edge entering a tree node is a tree edge, but if an edge enters a reticulate node it may be denoted a tree edge or a reticulate edge; we only require that at least one edge entering any given reticulate node be designated as a reticulate edge. This flexibility in whether we require both or only one edge entering a reticulate node to be designated as reticulate allows us to model different types of reticulation (e.g., in a hybridization network, both nodes entering a hybrid node would be reticulate, whereas in a “tree-based” network (Francis and Steel, 2015), one edge would need to be considered a tree edge).

Given these definitions, it is easy to see that a phylogenetic tree is a special case of a phylogenetic network, where there are no reticulate nodes, and hence also no reticulate edges.

##### Definition 1.

*If N is a phylogenetic network, then its unrooted version is obtained by ignoring the orientation of the edges and suppressing the root. The cycles in the unrooted version thus define a set of nodes in the rooted version that we refer to as* **cycles in the rooted network**. *A phylogenetic network N is said to be* **level-1** *phylogenetic network (or equivalently a* **galled tree**) *if no two cycles in the network share any nodes in common. If* γ *is a cycle in a level-1 network N and we restrict the nodes of N to those in* γ, *then a node in the restriction to* γ *with outdegree* 2 *(and hence indegree* 0) *is called a* **root** *of* γ, *and a node in the restriction with indegree* 2 *(and hence outdegree* 0) *is called a bottom of* γ.

Note that in Gusfield et al. (2003), the root of a cycle is referred to as a “coalescent node” and the bottom of a cycle is referred to as a “recombination node”. Using these definitions, the network shown in Figure 2 is a level-1 phylogenetic network that has a single cycle. The (unique) root of that cycle is *a* and the (unique) bottom of that cycle is *d*.

**Figure 2:**
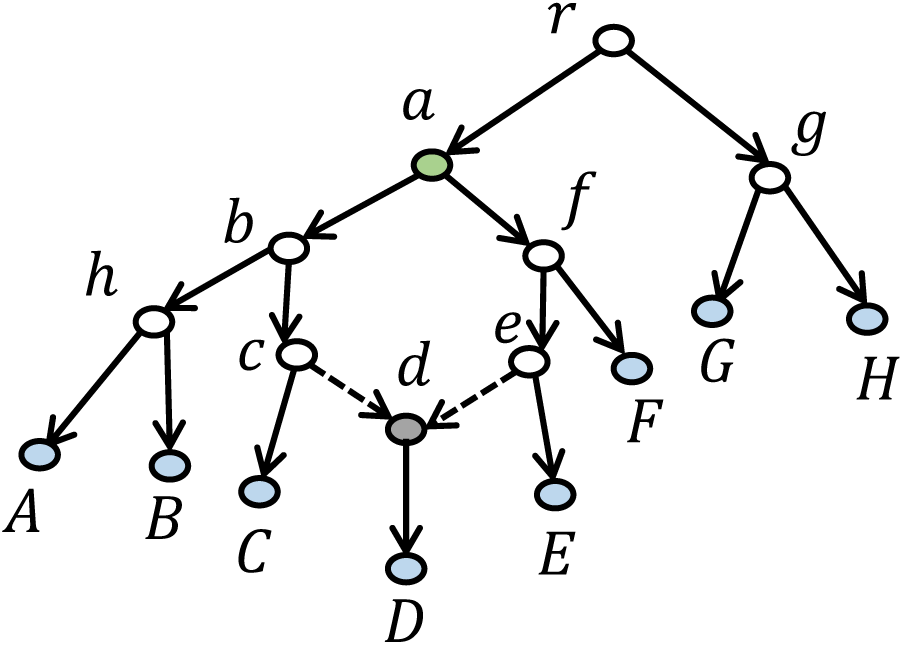
Level-1 phylogenetic network with one cycle of size six. Node *a* is the root of the cycle and node *d* is a reticulate node and is the bottom of the cycle; the edges entering *d* are both dashed, indicating that they are reticulate edges.

##### Observation 1.

*Suppose N is a level-1 network, and that when N is considered as an undirected graph, it has a cycle* γ. *Then there is a unique node y* ∈ γ *that has indegree* 0 *in the restriction of N to* γ *and a unique node x* ∈ γ *that has outdegree* 0 *in the restriction of N to* γ. *Furthermore, there are exactly two paths from y to x in N, and these two paths intersect only at x and y*.

*Proof.* This is a proof by contradiction.

Since γ is in a level-1 network, it follows that γ is chordless (i.e., a simple induced cycle), as otherwise there are at least two cycles in *N* that share vertices, contradicting that *N* is level-1. Now, since *N* does not have any directed cycles, there must be at least one node in γ that has indegree 0 and at least one node in γ that has outdegree 0, when considering only the subgraph of *N* induced by γ.

Suppose the root *r* of *N* is in γ, and suppose there is a second node *v* of indegree 0 in γ. Then at least one node on the path from *r* to *v* is not in γ, and so we have two cycles in *N* that are not node-disjoint.

On the other hand, suppose the root *r* is not in γ but that there are two nodes, *v* and *w* in γ that have indegree 0. If there is a directed path *P* from *v* to *w* or vice-versa, then again at least one node on this path is not in γ and so *N* has at least two cycles that are not disjoint. So, suppose *v* and *w* are in γ and have indegree 0 but there is no directed path from one to the other. Consider paths from the root of *N* to *v* and to *w*. Each of these paths contains at least one node that is not in γ, which means that *N* contains at least two cycles that are not node-disjoint. Therefore, there is exactly one node *y* in γ that has indegree 0 when restricting the attention to just γ. Since γ is a cycle, *y* has outdegree 2. We write the cycle in a circular ordering *y, v*_1_*, v*_2_*, . . ., v_k_, y*. Since *y* has indegree 0 it follows that the edges are oriented *y* → *v*_1_ and *v_k_* ← *y*. Hence there must be at least one node *x* in this cycle that has indegree 2. If there are two such nodes in the cycle, let the other be *x*′. We can write γ as the union of three paths, *P_y,x_* (the directed path from *y* to *x*), *P_y,x_′* (the directed path *y* to *x*′), and *P_x,x_′*, the path between *x* and *x*′. Since *x* and *x*′ both have indegree 2, it follows that this path contains at least one node that has indegree 0. But this path does not contain *y*, and so this means γ has two nodes of indegree 0, which we have already established is not the case. Therefore, γ has a unique node of indegree 0 and a unique node of outdegree 0.

It is easy to see that node *y* is the unique root of γ and *x* is the unique bottom of γ, according to Definition 1.

Finally, every simple induced cycle is formed of exactly two paths (*P*_1_ and *P*_2_) from its root *y* to its bottom *x*, and they intersect only at *x* and *y*. We now establish that there is no third path, *P*_3_ in *N* from *y* to *x*.

If there were a third path *P*_3_, then it must have at least one node that is not in γ. Hence, the subgraph obtained by picking *P*_1_ and the reversal of *P*_3_ forms a cycle that is different from *P*_1_ (and also different from *P*_2_, for that matter), and yet shares vertices *y* and *x*. Hence, *N* contains at least two cycles that intersect, contradicting that *N* is level-1.

##### Definition 2.

*Let* γ *be a cycle with at least three nodes in a rooted level-1 network N and let a and b be the root and bottom of* γ, *respectively. We will say* γ *is* **one-sided** *if there is an edge from a to b, and otherwise we say* γ *is two-sided*.

Note that the cycle in the network in Figure 1 has three cycles, where two are two-sided and one is one-sided.

##### Definition 3.

*For a rooted tree T with leafset S, given a node v in T, the set of leaves below v is referred to both as the clade at v and the cluster at v. The set of all such clades of a tree T is denoted by Clades(T)*.

It is well known that each rooted tree *T* is defined by its set of clades, and the tree can be constructed in linear time from the set (Gusfield, 1991).

A similar representation can be established for unrooted trees, but using bipartitions instead of clades, as we now describe.

##### Definition 4.

*Given an unrooted tree T with leafset S, if we delete an edge from T we partition the leaves into two sets A and B, and thus derive the bipartition A*|*B. The set of all such bipartitions for an unrooted tree with leafset S is denoted by Bip*(*T*). *Furthermore, if leaves a*_1_*, a*_2_ *are in A and b*_1_*, b*_2_ *are in B, then T induces the quartet tree a*_1_*a*_2_|*b*_1_*b*_2_*. The set of all such quartet trees of T is denoted by Q*(*T*)*. An equivalent definition for a*_1_*a*_2_|*b*_1_*b*_2_ *to be a quartet tree in Q*(*T*) *is that there are two node-disjoint paths in T, one connecting a*_1_ *to a*_2_ *and the other connecting b*_1_ *to b*_2_.

It is also well known that each tree *T* is uniquely defined by its set *Bip*(*T*), and can be constructed from *Bip*(*T*) in linear time (Gusfield, 1991). It is also true that the tree *T* can be constructed from *Q*(*T*) in polynomial time (Warnow, 2017).

##### Definition 5.

*Let N be a rooted phylogenetic network. If we delete one incoming edge for each reticulate node in N, we obtain a rooted tree T that we say is a tree* **contained in** *N. The set of all such trees is denoted by ᶯ_N_*.

As an example, there are exactly two trees in *𝒯_N_* for the network *N* given in Figure 2. We continue with some additional definitions based on *𝒯_N_* :

##### Definition 6.

*Let N be a rooted phylogenetic network, and recall that any tree T ∈ 𝒯_N_ is said to be a tree contained in N. We make the following definitions:*

- 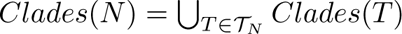
- 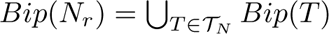
- 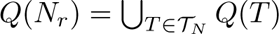

*Thus, Clades*(*N*) *is the set of clades of all the trees contained in N, Bip*(*N_r_*) *is the set of bipartitions of the trees contained in N, and Q*(*N_r_*) *is the set of quartet trees of trees contained in N. We also note that Clades*(*N*) *is sometimes referred to as the set of* soft-wired clusters *of the network N (Van Iersel et al., 2010)*.

We use the subscript *r* to emphasize that the sets are derived from trees contained in the rooted network *N*, and this very definitely depends on how *N* is rooted. We make this distinction explicit because in some studies (e.g., Gambette et al. (2012)), the set *Q*(*N*) of quartet trees is defined and used, and this set is different from *Q*(*N_r_*), as we now show.

##### Definition 7.

*Let N be a rooted phylogenetic network. Then the set Q*(*N*) *contains any quartet tree ab*|*cd (where a, b, c, d are leaves in N) so that N contains two node-disjoint paths P_ab_ and P_cd_, where P_ab_ connects a and b and P_cd_ connects c and d*.

Although trivially it follows that *Q*(*N_r_*) ⊆ *Q*(*N*), the two sets may not be identical, as we now show. Consider the network in Figure 2 and the leaf set *A, C, E, F* . This network has only one reticulate node, *d*, and there are two trees in *𝒯_N_*, depending on which edge entering *d* is discarded. However, both of those trees induce the same quartet tree on *A, C, E, F*, which is *AC*|*EF* . Thus, *Q*(*N_r_*) has only one quartet tree on *A, C, E, F*, which is *AC*|*EF* . By definition, *Q*(*N*) will contain that quartet tree, but we can see that it also contains *AF* |*CE*: there is a path connecting *A* and *F* (i.e., *A, h, b, a, f, F*) that is node-disjoint from a path from *C* to *E* (i.e., *C, c, d, e, E*).

We now further explore the question of when *Q*(*N*) and *Q*(*N_r_*) can differ. We begin with some definitions.

##### Definition 8.

*Let N be a level-1 phylogenetic network and let v be a node on a cycle* γ *in N and w a neighbor of v that is not in* γ; *therefore v is not the root of N. Note that* (*v, w*) *is a cut edge for N. Any leaf x in N for which the path to v passes through w is said to attach to* γ *at v. Each such leaf is also said to label the node it attaches to, but when we label a node v in a cycle, we arbitrarily pick one leaf that attaches to v for this labeling. Note that every leaf in N attaches to exactly one node in* γ, *and that the root of* γ *can be labeled by a leaf using this technique if and only if it is not also the root of N*.

Note that according to this definition, leaves *A* and *B* both attach to the cycle in Figure 2 at node *b* and so either leaf could be used to label node *b*. Note also that leaves *G* and *H* both attach to the root of the cycle. Finally, only one leaf attaches to the cycle at the other nodes. Note also that every leaf in the network attaches to exactly one node of the cycle.

We now prove that *Q*(*N*) and *Q*(*N_r_*) are always different whenever the network *N* is level-1 and has at least one cycle of length at least five.

##### Theorem 1.

*Let N be a rooted level-1 network. Then Q*(*N*) = *Q*(*N_r_*) *if all cycles are of length at most four, and otherwise Q*(*N_r_*) ⊊ *Q*(*N*) *(i.e., Q*(*N*) *contains Q*(*N_r_*) *as a proper subset)*.

*Proof.* If *N* is a level-1 phylogenetic network, it is easy to see that *Q*(*N*) = *Q*(*N_r_*) if all cycles are of length at most four.

Now consider the case where *N* is a level-1 phylogenetic network and has at least one cycle, γ, of length at least five. Let *r* denote the root of the cycle, and let *x* denote the bottom node of the cycle; all other nodes are “internal” nodes for γ. We label all the nodes of γ using leaves, according to the technique in Definition 8. We use a case analysis, based on whether γ is one-sided or two-sided.

The first case is where γ is two-sided. Since γ has at least five nodes, at least one side has two internal nodes, and we let the first two such internal nodes on that side (i.e. the ones closest to *r*) be labelled *a*_1_*, a*_2_. We let the first internal node on the other side (the one closest to *r*) be labelled *b*. Note that *r, a*_1_|*b, a*_2_ ∈ *Q*(*N*) \ *Q*(*N_r_*): there is an edge connecting the nodes labelled *r* and *a*_1_ a disjoint undirected path connecting the nodes labelled *b* to *a*_2_ that passes through the bottom of γ but the latter path is not a directed path in any tree in *N* .

The second case is where γ is one-sided, in which case it has at least three internal nodes on the same side. Using the same labeling technique of the internal nodes as in the previous case, let *a, b, c* be the first three nodes below the root, so that one side is the path *r, a, b, c, . . ., x*. It is easy to see that *ab*|*rc* ∈ *Q*(*N*)\ *Q*(*N_r_*): there is an edge connecting the nodes labeled *a* and *b* and a disjoint undirected path connecting the nodes labeled *r* and *c* that passes through the bottom of γ but the latter path is not a directed path in any tree in *N* .

Therefore, for any non-tree level-1 network *N* that has at least one cycle of length at least five, *Q*(*N*) ≠ *Q*(*N_r_*); moreover, every cycle of length at least five in the network contributes at least one quartet tree to *Q*(*N*) \ *Q*(*N_r_*).

### 2.2 Basic concepts about SNPs

In this paper we define SNPs to be two-state characters that have evolved down some tree *T* contained in the given level-1 phylogenetic network *N* under the infinite sites assumption. What this means is that the character evolves down *T* and changes state in exactly one edge of *T* . This is also described by saying the character evolves without homoplasy.

Note that if we are given a SNP and we know the ancestral state, then the set of leaves that exhibit the derived state will form a clade in some tree contained in the network. Thus, if we know the ancestral state for every SNP and we have enough SNPs, we can potentially derive *Clades*(*N*) and *Triplets*(*N*). Similarly, if we are given a SNP but we do not know the ancestral state, we can nevertheless infer a bipartition (i.e., split the leaf set into two parts: those leaves exhibiting the first state and those leaves exhibiting the second state). That bipartition will be in *Bip*(*N_r_*). Hence, if we are given enough SNPs, then even without knowing the ancestral state we can potentially obtain *Bip*(*N_r_*), and from this set we can obtain *Q*(*N_r_*).

## 3 Constructing Networks from SNPs

### 3.1 Overview

As described earlier, rooted level-1 phylogenetic networks can be constructed from their set of clades or rooted triplet trees, and hence also from SNPs with known ancestral state, provided that the SNPs cover the clades of the network. However, when the SNPs are given without known ancestral state, then clades and rooted triplet trees cannot be inferred, making the problem of inferring the network topology much more challenging. However, in this section we will show that the inference of the correct unrooted topology of a level-1 network *N* from SNPs without known ancestral state is guaranteed, provided that the network *N* has no cycles of size less than five and we have enough SNPs. To formalize this concept, we make the following definition

#### Definition 9.

*Let N be a level-1 phylogenetic network. We will say that a set of SNPs covers Bip*(*N_r_*) *if for every bipartition in Bip*(*N_r_*), *there is a SNP that produces that bipartition. Note that if a set of SNPs covers Bip*(*N_r_*) *then it also covers (in the same sense) Q*(*N_r_*).

Two prior algorithms, both described earlier, are potentially useful for constructing unrooted level-1 phylogenetic networks. The first is the polynomial time algorithm from Gusfield (2005) for constructing unrooted level-1 phylogenetic networks from SNPs, when the SNPs are given without an ancestral state and the recombination is unconstrained. We refer to this as the Gusfield-Construct-Unrooted algorithm, to credit the developer. The second is the polynomial time algorithm for constructing unrooted level-1 phylogenetic networks from quartet trees presented in Gambette et al. (2012). We refer to this method as the GBP algorithm to credit the three developers Gambette, Berry, and Paul.

We now address the problem of correctly constructing the unrooted topology of a level-1 phylogenetic network *N* from SNPs, when the ancestral state is unknown and all cycles are of size at least five. Since SNPs can be used to construct bipartitions or quartet trees on the leafset, this is closely related to the question of whether we can reconstruct the unrooted topology of a level-1 network *N* from *Bip*(*N_r_*) or *Q*(*N_r_*). A major contribution in this section is a proof that the GBP algorithm will fail to reconstruct *N* when given *Q*(*N_r_*). We also present a new quartet-based algorithm, CUPNS (Constructing Unrooted Phylogenetic Networks from SNPs), and prove it correctly reconstructs any level-1 phylogenetic network *N* when given SNPs that cover *Q*(*N_r_*), as long as all cycles are of length at least five. As a consequence, this establishes that *Q*(*N_r_*) is sufficient to define the unrooted topology of a level-1 network *N* where all cycles are of length at least five. Hence, this section also establishes that Gusfield-Construct-Unrooted correctly recovers the unrooted topology of *N* when given SNPs without known ancestral state and allowing for multiple crossovers, as long as they cover *Q*(*N_r_*) and *N* has no cycles of length smaller than five.

### 3.2 The GBP algorithm fails on *Q*(*N_r_*)

We begin by showing that the GBP algorithm fails on every level-1 network that contains at least one cycle of length at least five. Here we present, at a high-level, the GBP algorithm, when given an input set of quartet trees *Q*.

The input to the GBP algorithm is a set *Q* of quartet trees on leafset *S*. GBP then has the following steps.

- Step 1 (SN-tree computation): The SN-tree is computed. This is the unique unrooted tree *T*, leaf-labelled by *S*, that is the maximally resolved tree that satisfies the following two properties: (1) If *ab*|*cd* ∈ *Q*(*T*) then *ab*|*cd* ∈ *Q*, and (2) If *ab*|*cd* ∈ *Q*(*T*) then *ac*|*bd* ∉ *Q*. In other words, every resolved quartet tree in *T* is in *Q*, and there is no quartet tree in *Q* that conflicts with a resolved quartet tree in *T* .
- Step 2 (Polytomy resolution): The polytomies (i.e., nodes of degree greater than three) in the SN-tree are expanded into cycles.

The SN-tree can be calculated in polynomial time by (arbitrarily) ordering the leaves, and then building the SN-tree incrementally. As we will show later, if *Q* = *Q*(*N*) or *Q* = *Q*(*N_r_*), then the SN-tree is identical topologically to taking *N* and collapsing every cycle to a single node.

For Step 2, where the polytomies are expanded into cycles, the polytomies are expanded independently, and so it suffices to describe how a single polytomy is handled. First, given a polytomy *v* with degree at least four, each neighbor *x* is labelled by a leaf *l* whose unique path in the SN-tree to *v* passes through *x*. Once each neighbor of *v* is labelled, the GBP algorithm constructs a circular ordering of the neighbors using the “node ordering algorithm”, which has the following logic. Given three neighbors of *v* labelled *a, b, c*, the GBP algorithm decides that these nodes appear consecutively and in that order *if and only if* there does not exist a neighbor of *v* labelled *f* such that *bf* |*ac* ∈ *Q*. Then, having analyzed all triplets of neighbors, GBP returns a circular ordering *σ* on the neighbors of *v* if and only if the information gathered is consistent with *σ*. If such a circular ordering is found, then the polytomy is replaced by the cycle defined by *σ*.

#### Theorem 2

(From Gambette et al. (2012)). *Let N be a level-1 phylogenetic network. Then GBP will return the unrooted topology of N if given Q*(*N*) *as input*.

We note that the proof of correctness for the GBP algorithm explicitly assumes that the input set *Q* is identical to *Q*(*N*). However, we will prove in Theorem 3 that GBP *will always* fail if the quartet tree set given as input to GBP is limited to *Q*(*N_r_*) and the network *N* has at least one cycle of length at least five.

#### Lemma 1.

*Let N be a rooted level-1 phylogenetic network in which all cycles have at least four nodes. Let Z be any set of four leaves. Then Q*(*N_r_*) *has exactly two quartet trees on Z if and only if there is a cycle* γ *in N such that all four nodes of Z attach to different nodes in* γ, *and at least one of these leaves labels the bottom node in* γ.

*Proof.* For any four leaves, there is always at least one quartet tree in *Q*(*N_r_*). So assume that for some set *Z*, there are at least two quartet trees in *Q*(*N_r_*). If there is no cycle γ for which the four leaves in *Z* attach to four different nodes, then the four leaves of *Z* split 2|2 across some cut edge of *N*, and so there is only one quartet tree in *Q*(*N_r_*) for *Z*. Hence, there is at least one cycle γ in which the four leaves of *Z* attach to four different nodes. Furthermore, there cannot be two such cycles, because then the network is not level-1.

Now suppose there is a single cycle γ in which the four leaves in *Z* attach to different nodes, and suppose that the bottom node is not one of the nodes to which they attach. In that case, there is only one quartet tree on *Z* in *Q*(*N_r_*), as the choice of edge to delete from the two entering the bottom node will not impact the tree on *Z*.

Hence, since there are at least two quartet trees in *Q*(*N_r_*) for *Z*, it must be that the four leaves attach to different nodes in γ and one of the leaves in *Z* attaches to the bottom node, which is labelled *x*. Suppose the other nodes in γ are labelled *u, v, w* as you move from *x* in clockwise order. The cycle can be seen as consisting of two paths between *x* and *w*, one (*P_left_*) moving on the left-hand side going clockwise from *x* and the other (*P_right_*) moving on the right-hand side, going counter-clockwise from *x*. There are two edges entering *x*, one on the left (i.e., the first edge in *P_left_*) and one on the right (i.e., the first edge in *P_right_*). If we remove the first edge in *P_left_*, we obtain the quartet tree *xw*|*uv* on *Z*, and if we remove the first edge in *P_right_* we obtain the quartet tree *xu*|*vw* on *Z*. Thus, both quartet trees exist in *Q*(*N_r_*), but the third quartet tree does not exist in *Q*(*N_r_*), if one of the leaves attaches to the bottom node.

Thus, there is only one case where there is more than one quartet tree on *Z*, and it is the case where all four leaves of *Z* attach to some cycle γ at different nodes and one of the leaves attaches to the bottom node of γ, and in this case there are exactly two quartet trees on *Z*, as claimed.

We continue with a lemma that we will need for Theorem 4.

#### Lemma 2.

*Let N be a rooted level-1 phylogenetic network in which all cycles are of length at least five. Then the SN-tree computed using Q*(*N_r_*) *is identical to the SN-tree computed using Q*(*N*).

*Proof.* Recall that the SN-tree computed in Step 1 of GBP, given quartet set *Q* as input, is the maximally resolved unrooted tree *T* such that (1) *Q*(*T*) ⊆ *Q* and (2) no quartet tree in *Q*(*T*) conflicts with any quartet tree in *Q*. Consider then the tree obtained by taking the unrooted topology for *N* and collapsing every cycle into a polytomy (i.e., contracting all the edges in the cycle). Clearly, this tree satisfies the properties (1) and (2) above, and is maximally resolved subject to satisfying (1) and (2), when *Q* = *Q*(*N*). Here we show that this tree also satisfies the two required properties and is maximally resolved subject to this when *Q* = *Q*(*N_r_*).

It is easy to see that the tree satisfies (1) and (2). If some refinement *T* ^∗^ also satisfies (1) and (2), then there is a polytomy *v* in *T*, some pair of neighbors of *v* (which we will call *v*_1_*, v*_2_) so that adding a new node *v*′ and making it adjacent to *v, v*_1_, and *v*_2_ satisfies (1) and (2). Equivalently, that means finding two leaves *a, b* that attach to different nodes (*v*_1_ and *v*_2_) in the cycle that was collapsed to this polytomy, so as adding an edge that separates *a, b* from the rest of the leaves labeling the neighbors of the polytomy does not violate the two conditions above.

Note that the bottom node of the cycle is labeled by some leaf, *x*. Recall that, by Lemma 1, any quartet involving *x* and three other nodes in the cycle has two quartet trees in *Q*(*N_r_*). If *x* ∈ {*a, b*}, then picking any two other leaves labeling different nodes in the cycle than those labeled by *a, b* produces a set of four leaves for which there are two quartet trees. That means that this refinement, *T* ^∗^, produces a quartet tree that conflicts with a quartet tree in *Q*(*N_r_*). If *x* ∉ {*a, b*}, then we pick *x* and any other leaf *y* labeling some other node in the cycle, to get the set of four leaves *a, b, x, y*. This set of four leaves also has two quartet trees in *Q*(*N_r_*), meaning that the refinement *T* ^∗^ produces a quartet tree that conflicts with a quartet tree in *Q*(*N_r_*). Thus, no matter how *a, b* is picked, the tree *T* ^∗^ we obtain violates property (2). Hence, no refinement of this tree will satisfy the two required properties, and thus the SN-trees defined using *Q*(*N*) and defined using *Q*(*N_r_*) will be the same.

#### Theorem 3.

*Let N be a level-1 phylogenetic network in which at least one cycle is of size at least five. Then the GBP algorithm applied to Q*(*N_r_*) *will fail to correctly construct the unrooted topology of N*.

*Proof.* We already know that the SN-tree given *Q*(*N*) is the same as the SN-tree given *Q*(*N_r_*) from Lemma 2. We now show that the expansion of the polytomies in the SN-tree will fail to be correct whenever *N* satisfies the conditions above and GBP is given *Q*(*N_r_*).

Let *N* be an arbitrary rooted level-1 phylogenetic network for which at least one cycle γ has length at least five. Let *r* be the root of γ and let *x* be the bottom of γ. We label all the nodes in γ using leaves, according to the technique given in Definition 8. The proof then proceeds in a case analysis based on γ.

The first case is where γ has at least one internal node on each side. Since γ has at least five nodes, at least one side has two internal nodes, and we let the first two such internal nodes on that side be labelled *a*_1_*, a*_2_. We let the first internal node on the other side be labelled *b*. Note that *r, a*_1_|*b, a*_2_ ∈ *Q*(*N*) \ *Q*(*N_r_*). Now, the SN-tree computed in the first stage of the GBP algorithm will replace the cycle γ by a polytomy, with neighbors of the polytomy labelled using the same labels as how we labelled the cycle. The GBP algorithm will determine that the nodes labelled *b, a*_1_*, a*_2_ appear consecutively in that order if and only if the input set of quartet trees does not contain a quartet tree of the form *a*_1_*, z*|*b, a*_2_. Since no such quartet tree exists in *Q*(*N_r_*), GBP will determine that nodes *b, a*_1_*, a*_2_ appear consecutively in that order, which is a mistake. Hence, GBP will fail to correctly resolve the polytomy for this cycle when given *Q*(*N_r_*).

The second case is where the cycle γ is one-sided, in which case there are at least three internal nodes on one side of the cycle. Let the first three internal nodes below the root of the cycle be *a*_1_*, a*_2_*, a*_3_. The SN-tree will contain a polytomy to represent this cycle, which will have neighbors labelled using the leaves as we have described. GBP will consider *r, a*_1_*, a*_3_ consecutive if and only if there is no leaf *z* such that *a*_1_*, z*|*r, a*_3_ ∈ *Q*(*N_r_*). But there is no such leaf *z*, and so GBP will infer that nodes *r, a*_1_*, a*_3_ appear consecutively and in that order, which means GBP will fail to correctly resolve the polytomy in the SN-tree.

Thus, GBP will fail to correctly resolve every polytomy in the SN-tree when given *Q*(*N_r_*) as input, where *N* is a level-1 phylogenetic network with at least one cycle of size at least five.

We have established that GBP fails on all inputs *Q*(*N_r_*) for any level-1 phylogenetic network *N* with at least one cycle of length at least five. In contrast, if the input to GBP were *Q*(*N*), GPB would not make these mistakes, and would correctly resolve every polytomy into a cycle. The reason for this distinction is that, as established in Theorem 1, *Q*(*N_r_*) is a proper subset of *Q*(*N*) whenever *N* has at least one cycle of length at least five. However, we demonstrated that the quartet trees in *Q*(*N*) \ *Q*(*N_r_*) are needed for GBP to correctly expand polytomies in the SN-tree into cycles.

### 3.3 Gusfield-Construct-Unrooted

We begin by describing the algorithm presented in Gusfield (2005) for constructing an unrooted level-1 phylogenetic network, when given a set of SNPs (without known ancestral state) that covers the bipartitions in *N*, and we call this the Gusfield-Construct-Unrooted algorithm.

First, Gusfield-Construct-Unrooted constructs the character incompatibility graph, where the SNPs (i.e., two-state characters) are the nodes in the graph and edges exist between two SNPs if they are incompatible. It then computes a tree on the leafset using the characters that are compatible with all the characters in the input (i.e., this is a perfect phylogeny for this subset of the characters). If this tree does not have any polytomies, then it returns the tree. Otherwise, it uses the SNPs to replace each of the polytomies by cycles. We provide a description of this technique that is equivalent to what is described in Gusfield (2005), but expressed so as to enable a comparison to the GBP algorithm.

For a single polytomy *v* with degree *d*, the algorithm first labels each neighbor *w* by a single leaf (drawn arbitrarily from the other side of *w*, so that the path from the leaf to *v* passes through *w*). Then, it finds a neighbor *x* of *v* so that there is a perfect phylogeny on the leaf set labelling all neighbors of *v* other than *x*. Under the assumption that the input set of SNPs covers *Bip*(*N_r_*), there will be a perfect phylogeny and it will be a path with two cherries. This path is then closed into a cycle by looking at where *x* appears in the SNPs. This expansion of each polytomy into a cycle is performed independently across the polytomies, and then the resultant level-1 network is returned.

Gusfield (2005) proved that Gusfield-Construct-Unrooted returns a level-1 network consistent with the input SNPs, provided that the SNPs covered *Bip*(*N_r_*), and does so in polynomial time, but uniqueness was not established.

Also provided in Gusfield et al. (2003), and discussed in Gusfield (2005), is an algorithm for the case where the SNPs are given with known ancestral state. The only difference between the algorithms (one for rooted and one for unrooted SNPs) is that the compatibility tree produced in the algorithm is rooted when the SNPs are rooted, and the resolution of polytomies for the rooted case is more straightforward. Moreover, Gusfield et al. (2003) provides a proof of a unique solution, when given rooted SNPs (provided they cover the clades of the level-1 network). We refer to this algorithm as Gusfield-Compute-Rooted.

### 3.4 CUPNS: A new method for constructing unrooted phylogenetic networks from SNPs

We now present the Constructing Unrooted Phylogenetic Networks from SNPs algorithm (short form: CUPNS). CUPNS is designed to construct the correct unrooted level-1 network from SNPs given without ancestral state information, but will only be guaranteed to return such a network when the SNPs cover *Bip*(*N_r_*) and *N* has no cycles of size less than five. CUPNS combines features of the GBP algorithm with the algorithm in Gusfield (2005), as we show.

CUPNS has the following three phases:

- Phase 1 (Compute *Q_SNP_*). We use the SNPs to produce a set of quartet trees (all the quartet trees defined by the SNPs), which we denote by *Q_SNP_* .
- Phase 2 (SN-tree calculation). We construct the SN-tree *T* on *Q_SNP_* .
- Phase 3 (Polytomy resolution). For each polytomy *v* in *T* :

– If *v* has degree four, we reject (inconsistent with the required condition that all cycles be of length at least five). Otherwise, we label the neighbors of *v* using leaves, using the same technique as for the GBP algorithm, and let *L*(*v*) denote the set of leaves labelling the neighbors of *v*.
– We examine all the quartet trees on *L*(*v*) to find a leaf *x* such that every quartet of leaves in *L*(*v*) that includes *x* has exactly two quartet trees in *Q_SNP_* . If there is no such leaf or more than one such leaf, we return FAIL, indicating that no level-1 phylogenetic network is consistent with the input. Otherwise, we let *x* be the unique leaf that has exactly two quartet trees in *Q_SNP_* .
– We remove *x* from *L*(*v*). We construct a perfect phylogeny for the remaining leaves in *L*(*v*), if it exists, using Gusfield (1991); if no perfect phylogeny exists, we return FAIL. If the perfect phylogeny exists, it will be a path with two cherries (pairs of leaves that are siblings). We use the quartet trees in *Q_SNP_* to determine how to add *x* and close the cycle.

Note that Phase 2 (constructing the SN-tree) is identical to the first phase of the GBP algorithm. Furthermore, Phase 3 (polytomy refinement) is very similar to the algorithm in Gusfield (2005) except that we find the unique bottom node *x* based on Lemma 1, which is our contribution. Thus, CUPNS has features from both the GBP algorithm and Gusfield (2005). Note that the algorithm can fail to return a network, but this can only occur during the third phase (polytomy resolution), and failures only occur if the input SNPs are inconsistent with the required conditions (i.e., the network is level-1, all cycles have at least five nodes, and the SNPs cover the bipartitions of the network). However, optionally, we may elect to return the SN-tree, or the partially modified version of the SN-tree prior to the failure. On the other hand, when the input is consistent with the required conditions, we can prove that CUPNS returns the true unrooted topology of the network.

#### Theorem 4.

*Let N be a rooted level-1 phylogenetic network in which all cycles are of length at least five. If we apply CUPNS to a set of SNPs that covers Q*(*N_r_*), *then we obtain the unrooted topology for N, and the bottom node for each cycle is also identified*.

*Proof.* By Lemma 2, the SN-tree computed using *Q*(*N_r_*) is identical to the SN-tree computed using *Q*(*N*). We now show that the second step where we replace the polytomies in the SN-tree by cycles returns the unrooted topology of *N*, when the SNPs cover *Q*(*N_r_*). Let *v* be a polytomy and *L*(*v*) be the set of leaves labeling the neighbors of *v*. Since every cycle in *N* has at least five nodes, for all sets of four leaves labeling different nodes in the cycle, where none of them label the bottom node, there is exactly one quartet tree in *Q*(*N_r_*) (Lemma 1). Furthermore, for any set of four leaves labeling different nodes in the cycle, if one of them labels the bottom node, there are two quartet trees in *Q*(*N_r_*) (Lemma 1). Hence, the algorithm correctly identifies the leaf *x* labeling the bottom node of each cycle.

The algorithm computes the set *L*(*v*)\{*x*} of leaves that label the remaining nodes of the cycle. It is easy to see (but see Gusfield (2005) for additional discussion and algorithmic details) that this set has a perfect phylogeny (i.e., a tree on which all the SNPs evolve without homoplasy), and since binary character perfect phylogenies are unique (Gusfield, 1991), the perfect phylogeny is the caterpillar tree with two cherries formed by deleting the edges incident with the bottom node from the cycle. As described in Gusfield (2005), there is also a unique way to add back in the leaf labeling the bottom node to create the cycle that is consistent with the quartet trees in *Q*(*N_r_*), and it can be computed efficiently. Hence, the algorithm produces the correct cycle for each polytomy in the SN-tree, establishing correctness.

Hence, we obtain the following theorem and corollary.

#### Theorem 5.

*Let N be a level-1 rooted phylogenetic network with all cycles of length at least five. Then, the set Q*(*N_r_*), *as well as the set Bip*(*N_r_*), *are each sufficient to define the unrooted topology of N*.

*Proof.* Let *N* be a level-1 phylogenetic network where all cycles are of length at least five. Recall that CUPNS operates by using the SNPs, which do not have a given ancestral state, to define quartet trees. Thus, if the SNPs cover the clades of *N*, it follows that CUPNS obtains every quartet tree in *Q*(*N_r_*). Now recall that Theorem 4 established that CUPNS will return the unrooted topology of *N* given a set of SNPs, without known ancestral state, provided that the input SNPs cover the clades of *N* . Hence it follows that *Q*(*N_r_*) is sufficient to define the unrooted topology of *N* . Note also that the set *Bip*(*N_r_*) of bipartitions of trees in *N* can be used to obtain *Q*(*N_r_*), thus establishing that *Bip*(*N_r_*) is sufficient to define the unrooted topology of *N*.

Gusfield (2005) established that Gusfield-Construct-Unrooted is guaranteed to return *an unrooted level-1 network* consistent with the input SNPs evolving down a rooted version of the output network without homoplasy, provided that the SNPs covered the clades of the network. However, Gusfield (2005) did not establish that there was only one possible unrooted network consistent with such an input. We now establish uniqueness is in fact the case.

#### Corollary 1.

*Let N be a level-1 rooted phylogenetic network with all cycles of length at least five*. Gusfield-Construct-Unrooted *will return the unrooted topology of N if the SNPs cover Bip*(*N_r_*)*. Furthermore,* Gusfield-Construct-Unrooted *runs in O*(*mn* + *n*^3^) *time, where N has n leaves and there are m SNPs*.

*Proof.* Theorem 5 establishes that any two level-1 networks that have the same set *Bip*(*N_r_*) of bipartitions from the trees they contain must be topologically identical. Since Gusfield (2005) established that the output unrooted network is consistent with each of the input SNPs evolving down some tree contained in the rooted network, this directly proves that the output of Gusfield-Construct-Unrooted must be the correct unrooted network topology–provided that the input SNPs cover *Bip*(*N_r_*). The runtime of Gusfield- Construct-Unrooted was proven in Gusfield (2005) to be *O*(*mn* + *n*^3^) time, where there are *m* SNPs and *n* taxa.

### 3.5 Summary

In this section, we addressed the question of how to construct the unrooted topology of a level-1 phylogenetic network *N* in which all cycles are of length at least five given a set of SNPs, without ancestral state information, that covers *Bip*(*N_r_*), under the assumption that the SNPs evolve down *N* under the infinite sites assumption. We proved that the GBP method, applied to the quartets produced by the SNPs (i.e., *Q*(*N_r_*)) will fail on every level-1 network that has at least one cycle of length at least five. However, we also presented two methods, CUPNS (the method we introduced) and Gusfield-Construct-Unrooted, each of which has a guarantee of returning the unrooted topology of *N* given such a set of SNPs. Fur- thermore, Gusfield-Construct-Unrooted runs in *O*(*mn* + *n*^3^) time where there are *m* SNPs and *n* leaves, while CUPNS has a step that requires it compute all the quartet trees, and so its runtime is higher than Gusfield-Construct-Unrooted. We summarize this discussion for constructing unrooted level-1 phylogenetic networks from SNPs in Table 1.

**Table 1:** Properties of algorithms for constructing unrooted level-1 networks from SNPs. The “required conditions” are that *N* is a level-1 network without cycles of size less than five, and that the SNPs cover *Bip*(*N_r_*). The runtimes for GBP-SNPs and CUPNS assume that it takes *O*(*mn*^4^) time to compute all the quartet trees from the input set of *m* SNPs, and that this dominates the total runtime.

| Method | Comments if required conditions hold | Runtime |
| --- | --- | --- |
| Gusfield-Construct-Unrooted | Guaranteed correct | $O(mn + n^3)$ |
| CUPNS | Guaranteed correct | $O(mn^4)$ |
| GBP-SNPs | Will fail on all non-tree networks | $O(mn^4)$ |

## 4 Model of SNP Evolution and Network Estimation

### 4.1 Preliminary discussion

In this section, we describe the stochastic model of nucleotide evolution down a level-1 phylogenetic network, and then show how we can estimate the model phylogenetic network from the alignment that is produced.

We will refer to the leaves of the phylogenetic network as “species”, but in some contexts (e.g., population genetics) they may be individuals rather than species.

Recall that in this paper SNPs, by definition, are assumed to exhibit two states and evolve under the infinite sites assumption (i.e., without homoplasy) down some tree contained in the level-1 network *N* . However, the sites within an alignment that are SNPs are not known. Therefore, in order to use a multiple sequence alignment to construct a phylogenetic network based on SNP analysis, the biologist must first decide which sites are likely to be SNPs, and then restrict the analysis to that subset of the sites.

Here we model this selection process by saying that the biologist has access to an “Oracle”, which tells them which sites are SNPs (i.e., exhibit two states and evolve under the infinite-sites assumption) and possibly also the ancestral state for the SNPs.

We describe a character evolution model that is based on DNA sequence evolution, with *i.i.d.* evolution across the sites. The model allows for homoplasy in the usual way, but has the Oracle identify the sites that evolved without any homoplasy, changing only once in the evolutionary history, and so exhibit two states. In other words, the Oracle identifies the SNPs from the alignment.

Given a level-1 phylogenetic network *N*, each character essentially “picks” a tree in *𝒯_N_* down which to evolve, and then evolves down that tree under a standard model involving substitutions of nucleotides by other nucleotides. Hence, to define the character evolution model, we need to define how each character “picks” the tree down which to evolve, and also the substitution model.

Here we describe the numeric parameters of the level-1 phylogenetic network *N* and how they govern character evolution.

### 4.2 Single site evolution model

The network numeric parameters and the constraints on these parameters are as follows:

- The state at the root is random and each of the four nucleotides has a strictly positive probability of appearing at the root.
- Every edge *e* entering a reticulate node has a transmission probability *κ*(*e*) with 0 *< κ*(*e*) *<* 1 and the transmission probabilities of the edges entering a reticulate node sum to 1.
- For a directed edge *e* = (*a, b*) and nucleotides *s, s*′, let *p*(*e*; *s, s*′) be the conditional probability of observing *s*′ at node *b* given that *s* is observed at node *a*. Then, for all tree edges *e*, *p*(*e*; *s, s*′) *>* 0 for all nucleotides *s≠ s*′, and, for all reticulate edges *e*, *p*(*e*; *s, s*′) = 0 for all nucleotides *s*≠ *s*′,

We now provide some comments on how these parameters influence the evolution of a single character down the network.

- Because the transmission probabilities of the edges entering a reticulate node must sum to 1, for each character, each reticulate node inherits from exactly one of its parents. Thus, each character evolves down a tree contained in *𝒯_N_* .
- Because characters evolve *i.i.d.*, different characters can be inherited from different parents.
- Because reticulate edges represent the transfer of character states at a specific time, such as in a lateral gene transfer, we have *p*(*e*; *s, s*) = 1 for a reticulate edge *e* = (*v, w*) and nucleotide *s*. This has the consequence that the state of character *c* at *v* is the same as its state at *w* where *c* is a character that is transmitting on *e*.

Overall, therefore, this character evolution model can be seen as enforcing that every character evolves down a single tree in *𝒯_N_*, with a probability that depends on the parameter values for *κ*(*e*) for all edges *e* entering reticulation nodes, but that every tree in *𝒯_N_* has strictly positive probability of being selected; once the tree is selected, the substitution process is specified by the substitution probabilities *p*(*e*; ·, ·) on the edges *e* in the tree it selected.

### 4.3 Estimating the rooted level-1 phylogenetic network

Although our major interest is in estimating unrooted level-1 networks from SNPs without known ancestral state, establishing statistical properties of methods for the corresponding problem of estimating rooted level-1 networks from SNPs with known ancestral state is also of interest. Hence, we begin with this case.

#### 4.3.1 Gusfield-Estimate-Rooted

We describe a method we call Gusfield-Estimate-Rooted to estimate the topology of a level-1 phylogenetic network *N* from a DNA alignment of *n* sequences and *k* sites, under the assumption that the sites have evolved down *N* . We begin with a definition we will need.

##### Definition 10.

*Let N be a level-1 phylogenetic network. A set of rooted SNPs is said to cover Clades*(*N*) *if and only if for every clade in Clades*(*N*) *there is at least one rooted SNP defining the same bipartition (with the nodes in the clade having the derived state)*.

We now describe the use of an algorithm from Gusfield (2005) for the estimation problem when the SNPs are given with known ancestral state.

- Stage 1: Given the *k* DNA characters (equivalently, a DNA alignment of *k* sites), the Oracle determines which of them are SNPs (i.e., evolved without homoplasy down some tree contained in the network and exhibit only two states) and identifies the ancestral state for each such SNP (thus producing rooted SNPs).
- Stage 2: Apply Gusfield-Construct Rooted Phylogenetic Network from Rooted SNPs to the rooted SNPs.

We call this two-step approach Gusfield - Statistical Estimation of Rooted Phylogenetic Net-work from Rooted SNPs (or more simply Gusfield-Estimate-Rooted), to emphasize that this is a statistical estimation approach. Note that this technique uses the input alignment in a two-stage process, where the first stage extracts the rooted SNPs, and these are then used to construct a level-1 phylogenetic network in the second stage.

#### 4.3.2 Statistical consistency, sample complexity, and runtime

##### Theorem 6.

*Let N be a level-1 phylogenetic network and suppose that characters evolve down N under the model described in Section 4. Suppose also that the Oracle does not make any mistakes, so that it correctly identifies all the SNPs and correctly specifies their ancestral states. Then, as long as all cycles in N are of length at least five*, Gusfield-Estimate-Rooted *will return the rooted network topology for N with probability converging to* 1 *as the number of sites increases (i.e., it is a statistically consistent estimator of the rooted network topology). Furthermore, the total runtime is O*(*kn* + *n*^3^)*, where k is the alignment length in the input and n is the number of leaves in the network*.

*Proof.* We begin by establishing statistical consistency. The proof of correctness for Gusfield-Construct-Rooted if the SNPs cover all the clades of the level-1 network is given in Gusfield et al. (2003). Hence, all we need to establish is that as the number *k* of characters increases, with probability converging to 1, the SNPs in the input will cover *Clades*(*N*) (see Definition 10).

For a given set *A* ∈ *Clades*(*N*) there is at least one tree *T* ∈ *𝒯_N_* such that *A* ∈ *Clades*(*T*). Note also that the probability of a character evolving down *T* is strictly positive, since 0 *< κ*(*e*) *<* 1 for all reticulate edges in *N*, and every edge is either a reticulate edge or a tree edge.

Since *A* is a clade in *T*, *v* = *lca_T_* (*A*) has outdegree two and therefore is a tree node in *N* . Hence there is a single edge *e* entering *v*, and since it is a tree edge it follows *p*(*e*; *s, s*′) *>* 0 for all nucleotides *s* ≠ *s*′. Now suppose that *c* is a character that evolves down *T* . It is easy to see that the probability that *c* changes on *e* and no other edge in *T* is strictly positive. Note also that if *c* evolves down *T* and changes only on edge *e*, then *c* is two-state and homoplasy-free, and hence is a SNP. Hence, the probability of *A* appearing as *clade*(*c*) for some SNP *c* is strictly positive.

Since we assume that the Oracle does not make any mistakes in identifying SNPs and their ancestral states, we can use the Oracle to identify the sites that are SNPs in an alignment and hence obtain a subset of the clades of *N* . Also, the previous argument shows that every clade in *N* has strictly positive probability of appearing as a clade for a SNP. Hence, as the number of sites increases, with probability converging to 1, the set of clades for the SNPs converges to *Clades*(*N*). Statistical consistency then follows, as noted above. The runtime analysis is straightforward. Gusfield et al. (2003) established that Gusfield-Construct- Rooted runs in *O*(*mn* + *n*^3^) time, where there are *m* SNPs and *n* taxa. The input alignment has *k* sites, of which *m* ≤ *k* are SNPs, and we assume that the Oracle takes *O*(1) time on each site. Hence, the runtime is *O*(*nk* + *n*^3^) time.

We now establish a bound on the sample complexity.

##### Theorem 7.

Let N be a level-1 phylogenetic network without cycles of length less than five. Assume that the Oracle identifies the SNPs and defines the ancestral state for every SNP. Suppose that 0 *< p*_on_ ≤ *p*(*e*; *s, s*) *and* 0 *< p*_off_ ≤ *p*(*e*; *s, s*′) for edges e in N that are not reticulation edges and states s ≠= *s*′*. Suppose also that* 0 *< κ*_1_ ≤ *κ*(*e*) ≤ *κ*_2_ *<* 1 *for all edges e entering reticulate nodes in N. For a given ɛ >* 0, Gusfield- Estimate-Rooted *returns N with probability at least* 1 − *ɛ when the number k of characters satisfies*

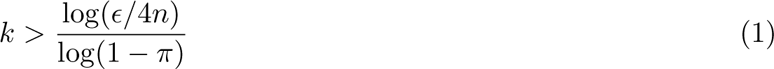

*for* 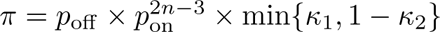

*Proof.* By the arguments provided in Theorem 6, Gusfield-Estimate-Rooted will return the rooted topology of *N* if the alignment has enough SNPs to cover all the clades of *N* . Therefore, we only need to compute the minimum number *k* of characters (sites) that guarantees that the SNPs cover *Clades*(*N*) with probability at least 1 − *ɛ*.

We begin with some observations that will be useful in this proof. Each edge in the network that is not in any cycle defines a unique bipartition, and hence clade (since we know the root state). Each edge in the network that is in a cycle, but is not incident to the bottom node of the cycle, defines two bipartitions (and hence two clades). Specifically, if γ is a cycle with bottom node *x*, then we let *X* denote the clade below the bottom node, and we refer to this as the bottom clade. Then any edge *e* in γ, with node *v* at the bottom end of *e*, denotes two clades: one containing *X* and one disjoint from *X*. We refer to these two as the *full clade* and *partial clade*, respectively, associated with edge *e* and hence also with node *v*. Furthermore, any clade *c* that is not the clade below the bottom node in a cycle or (in the case of a one-sided cycle) the clade below the root of the cycle is associated with a unique edge (and hence a unique node at the bottom of the edge). The clade that is below the bottom node of a cycle is associated with all three edges incident with the bottom node of the cycle.

The probability of the event that a clade *A* appears for the network as input from a given character is of the general form ∑_*T*_ *P*(*ε*(*T*) × *P*(*ε*(*A, T*) | *ε*(*T*)), where *ε*(*T*) is the event that the character evolves down the tree *T*, E(*A, T*) is the event that the character changes state on the edge in *T* associated with the clade *A*, and the sum is over trees *T* belonging to some suitable subset of *𝒯_N_*. The first probability in the sum is a product over the transmission probabilities *κ*(*e*) for edges *e* that enter nodes in *T* that were reticulate nodes in *N* . If the sum of these products is over all of *𝒯_N_* then it is, of course, 1.

Consider the conditional probability *P* (E(*A, T*) | E(*T*)) for a given tree *T* . Denote the distribution of the character at the root node by *ρ*. Let *e* be the edge in *T* associated with the clade *A*. The conditional probability we seek is

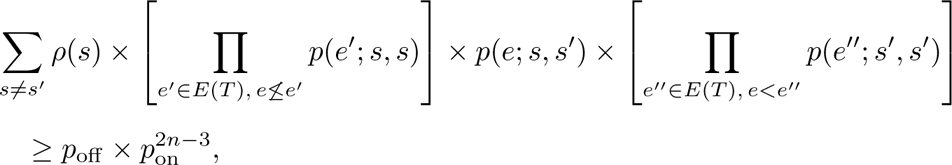

where {*e*′′ ∈ *E*(*T*) : *e < e*′′} are the edges in the tree *T* that are strictly below the edge *e* and {*e*′

∈ *E*(*T*)*, e* ≤*e*′} is the complement in *E*(*T*) of {*e*′′ ∈ *E*(*T*) : *e < e*′′} ∪ {*e*}.

We analyze the following cases: (1) clades associated with edges that are not in any cycle, (2) the partial clade associated with an edge in a cycle, and (3) the full clade associated with an edge in a cycle.

- **Case 1: The clade** *A* **is associated with an edge** *e_A_* **that is not in any cycle.** Since *e_A_*is not in any cycle, such an *e_A_* is an edge in every tree contained in *N* . Hence, when there is a substitution on edge *e_A_* and no other edge in the network, clade *A* appears in the input, no matter which tree the character evolves down. It follows from the discussion above that the probability we seek is at least 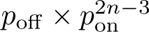
- **Case 2: The clade** *A* **is a partial clade associated with an edge in a cycle.** Let *X* be the bottom clade (i.e., the clade associated with the bottom node) and let *A* and *A* ∪ *X* be the partial clade and full clades associated with the node *v_A_* and the edge *e_A_*. Let *e_r_* be the edge entering the bottom node that is on the same side of the cycle as *v_A_*; note that *e*≠ *e_A_*, since *v_A_* is not the bottom node. The probability of transmission on edge *e_r_* is *κ*(*e_r_*). When a character only changes state on the edge *e_A_*and does not transmit on the reticulation edge *e_r_*, clade *A* appears in the data; and when the character transmits on *e_r_* then clade *A* ∪ *X* appears in the input. Now we must restrict our sum to trees *T* that do not transmit on edge *e_r_*. The sum of the corresponding set of products is then 1 − *κ*(*e_r_*). The conditional probability that clade *A* appears in the input given that we evolve down *T* is bounded below by 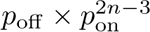. Thus, the probability we seek is at least 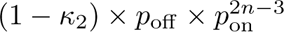.
- **Case 3: The clade** *A* **is a full clade associated with an edge in a cycle.** Let *T* be a tree in the network down which a character evolves that yields clade *A* ∪ *X*. Hence *T* must include the edge *e_r_*. The probability that clade *A* ∪ *X* appears in the input given that we evolve down *T* is at least 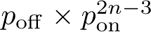. Note that we must transmit on edge *e_r_*. Thus, the probability we seek is at least 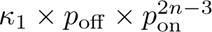.

Write 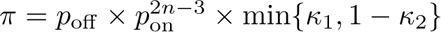 for the lower bound we have obtained for the probability that an arbitrary given clade appears for a given character. It follows that the probability that an arbitrary given clade appears for at least one of *k* characters is at least 1 − (1 − *π*)*^k^*. By Boole’s inequality (sometimes known as the union bound), an upper bound on the probability that some clade is missing from the list of clades for *k* characters is *c*(*N*)(1 − *π*)*^k^* ≤ 4*n*(1 − *π*)*^k^* where *c*(*N*) is the number of clades for the network *N* and the upper bound *c*(*N*) ≤ 4*n* is a consequence of Lemma 8.8.1 in Gusfield (2014). A lower bound on the probability that all clades are present in the list of clades for *k* characters is thus 1 − 4*n*(1 − *π*)*^k^*. This last quantity is greater than 1 − *ɛ* if and only if *k* is greater than the quantity on the right-hand side in the statement of the theorem.

### 4.4 Estimating the Unrooted Level-1 Phylogenetic Network

Here we wish to estimate the unrooted topology of the level-1 phylogenetic network, and so we do not need to know the ancestral state for each SNP. For the case where we do not have the ancestral state, GBP does not have theoretical guarantees for estimating the network from SNPs, but by Theorem 4 and Corollary 1, both CUPNS and Gusfield-Construct-Unrooted are guaranteed to return *N* if the SNPs cover *Bip*(*N_r_*) (and hence cover *Q*(*N_r_*)) and the network does not have cycles with length less than five. This is summarized in Table 1.

Therefore, we now describe the use of Gusfield-Construct-Unrooted for *statistical estimation* of the unrooted phylogenetic network from a DNA multiple sequence alignment.

#### 4.4.1 Gusfield-Estimate-Unrooted

- Step 1: Given the *k* DNA characters, the Oracle determines which of them are SNPs (i.e., evolved without homoplasy down some tree contained in the network and exhibit only two states).
- Step 2: Apply Gusfield-Construct Unrooted Phylogenetic Network from Unrooted SNPs to the SNPs.

We call this two-stage approach as Gusfield- Statistical Estimation of Unrooted Phylogenetic Network from Unrooted SNPs, or more simply as Gusfield-Estimate-Unrooted, to emphasize that this is a statistical estimation approach that operates on a given multiple sequence alignment.

#### 4.4.2 Statistical Consistency, Sample Complexity, and Runtime

##### Theorem 8.

*Let N be a level-1 phylogenetic network with n leaves, and assume all cycles have at least five nodes. Assume that character evolution down N is under the model from Section 4. If the Oracle specifies the SNPs but not the ancestral state, then as the number k of characters increases,* Gusfield-Estimate-Unrooted *will return the unrooted topology of N with probability converging to* 1 *(i.e. it is a statistically consistent estimator of the unrooted network topology). Furthermore, the total runtime is O*(*kn* + *n*^3^), *where k is the alignment length in the input*.

*Proof.* The correctness and runtime of Gusfield-Construct-Unrooted is given in Corollary 1 (see also Gusfield (2005)). The rest of the proof of consistency and runtime analysis follows as for the rooted case.

##### Theorem 9.

*Let N be a level-1 phylogenetic network without cycles of length less than five. Assume that the Oracle identifies the SNPs but does not define the ancestral state for every SNP. Suppose that* 0 < *p*_on_ ≤ *p*(*e*; *s, s*) *and* 0 < *p*_off_ ≤ *p*(*e*; *s, s*′) *for edges e in N that are not reticulation edges and states s≠s*′. *Suppose also that* 0 < *κ*_1_ ≤ *κ*(*e*) ≤ *κ*_2_ < 1 *for all edges e entering reticulate nodes in N. For a given* ɛ > 0, Gusfield-Estimate-Unrooted *returns N with probability at least* 1 − *ɛ when the number k of characters satisfies*

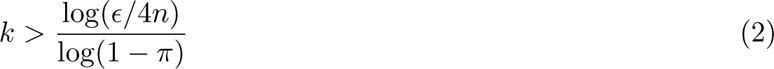

*for* 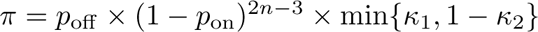

*Proof.* The argument for the sample complexity analysis for the unrooted case is identical to the rooted case, as we now show. Note that Theorem 8 establishes that Gusfield-Estimate-Unrooted will return the correct unrooted network *N* if the input alignment has SNPs that cover *Bip*(*N_r_*); this is the same set of SNPs that would cover the clades in *N* . Hence, the argument provided for the rooted SNP case can be modified for use in the unrooted SNP case.

## 5 Limitations and Extensions

In this section, we discuss the limitations of the theory we have established, as well as extensions to the case of multi-state characters. For limitations, we specifically consider whether CUPNS and the algorithm for constructing phylogenetic networks from SNPs without known ancestral state from Gusfield (2005) (i.e., Gusfield-Construct-Unrooted) are robust to various violations of the underlying assumptions (e.g., the true network has no small cycles, the Oracle does not make any mistakes, and there are enough SNPs to cover the bipartitions of the network).

### 5.1 Outcomes when the network has small cycles

In this paper we have always required that the network cycles be of length at least five. It is well known that rooted phylogenetic networks with cycles of length four are not identifiable from their clades or rooted triplet trees, and we provide an example of three networks with the same clades and triplet trees in Figure 3. The networks in this figure also have the same quartet trees, thus providing an example where the quartet trees in *Q*(*N_r_*) are insufficient to define the unrooted topology of *N* due to having four-node cycles.

**Figure 3:**
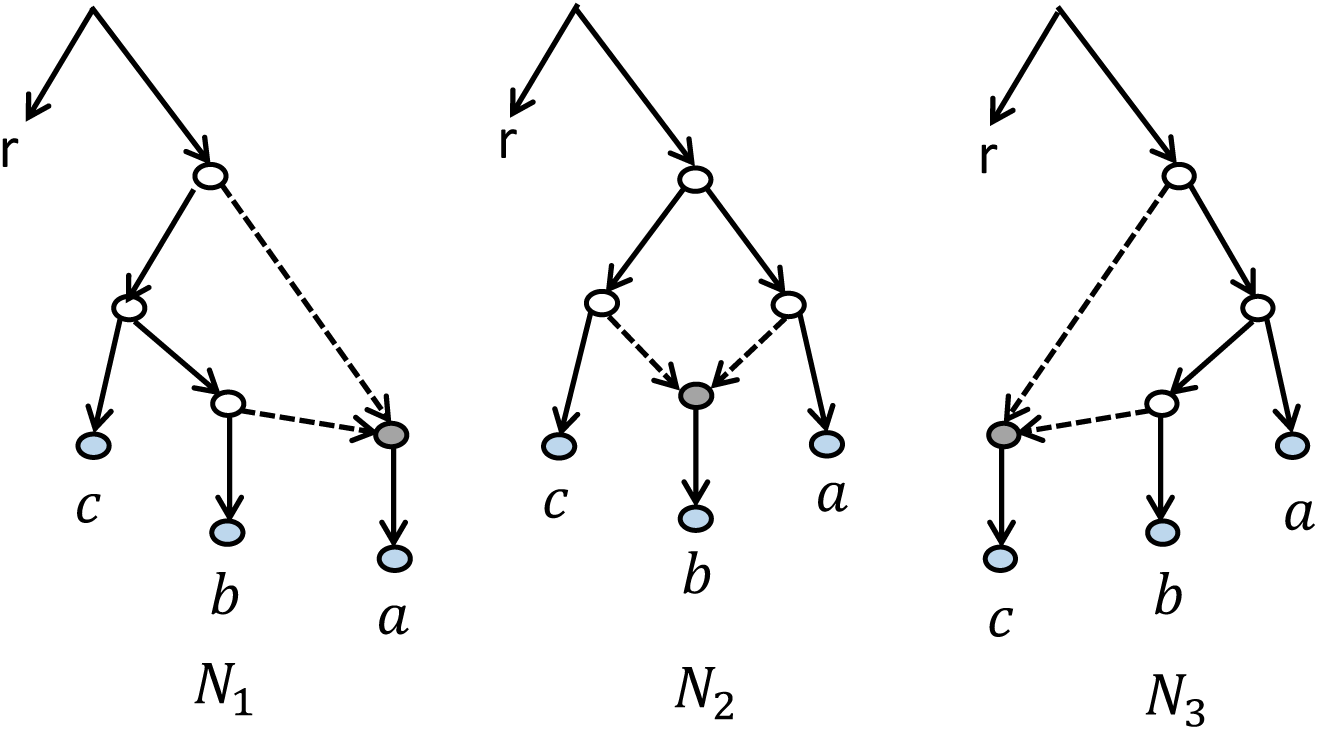
Three rooted level-1 networks on the same set of four leaves that have the same clade set and the same set of quartet trees induced by trees contained in the network, i.e., *ab cr* and *bc ar*. This establishes that rooted networks with four-node cycles are not identifiable from their clades or rooted triplet trees and their unrooted topologies are not identifiable from their quartet trees.

### 5.2 Outcomes when the SNPs are insufficient

The theoretical results we have established assume that the SNPs cover the network bipartitions. Here we examine the consequences when the SNPs cover some but not all of the network bipartitions. As we will show, if there are not enough SNPs to cover the clades, the result obtained can fail to be the correct network.

Recall that a rooted phylogenetic network *N* with at least one cycle has at least two trees in *𝒯_N_*. The first set of cases we describe have to do with the situation where the input SNPs are pairwise compatible.

The simplest case, and the most problematic, happens when, for example, each of the SNPs changes only on a cut edge of the network but the network has at least one cycle. This situation produces a set of compatible bipartitions, thus consistent with a single non-binary tree that is either identical to or a contraction of the SN-tree we would have obtained if we had SNPs defining all the bipartitions of the network. However, there is no information on how to resolve this tree, which would have at least one polytomy.

A less extreme version is when the SNPs all evolve down the same tree *T* ; here too the bipartitions produced by the SNPs are compatible, and the set of compatible bipartitions will define a single minimally resolved tree *T* ′ that will either be equal to *T* or a contraction of *T* . If every internal edge of *T* has at least one SNP changing on it, then *T* = *T* ′ is guaranteed. This case is somewhat more problematic, because there is no reason to suspect that we are missing information, and we might just return the tree as the “true network”, but it would not be correct.

Another version of this problem is when the SNPs evolve down different trees, but they do not change state on the right edges, with the result that the set of bipartitions they define may still be compatible but insufficient to define the network. For example, in the network given in Figure 2, there are two trees in the network (that differ only in the placement of *D*), and some SNPs might evolve down the first tree and others down the second tree. But if there are no changes on any edges below node *a*, for example, then there is no way to know that the true network is not a tree, since the SNPs are compatible with a tree phylogeny.

More generally, without some evidence that there are incompatible SNPs, no method can recover the true network. The best that can be obtained is that the method will return the minimally resolved tree compatible with all the SNPs.

The second set of cases we describe are where the input SNPs define a set of bipartitions that is incompatible, and so provides evidence of at least one cycle, but where there is not enough information to determine the correct network.

Consider, for example, the level-1 phylogenetic network in Figure 2. This network has one cycle. Suppose for every edge in this network other than (*a, f*), there is at least one SNP that changes on the edge. Suppose in particular that there is a SNP that changes on edge (*f, e*) and on no other edge and that uses the reticulate edge (*e, d*); this SNP produces the clade {*D, E*}. Suppose that there is a SNP that changes on edge (*b, c*) and on no other edge and that uses the reticulate edge (*c, d*). This SNP produces the clade {*C, D*}. If we know the ancestral states for the SNPs, then the clades we obtain from these two SNPs are incompatible, because {*C, D*} and {*D, E*} have a non-empty intersection and neither contains the other. If we do not know the ancestral states for the SNPs, then we get bipartitions instead of clades, but here too the bipartitions corresponding to these two clades are incompatible. Thus, whether we know the ancestral state for all the SNPs or not, the set of clades or bipartitions we obtain is incompatible, thus providing direct evidence that the true phylogenetic network has at least one cycle.

Consider the consequence of not having any SNP change on edge (*a, f*). This means there is no SNP providing evidence of the clade {*D, E, F* } (or equivalently the bipartition *A, B, C, G, H*|*D, E, F*). Since we assume there is a SNP changing on the edge (*r, a*) and on no other edge, we do have evidence for the bipartition *A, B, C, D, E, F* |*G, H* (or the clade {*A, B, C, D, E, F* } if we know the ancestral state). But note also that if we move *F* from its position in *N* to some other position we could produce a new network that is compatible with the SNPs. For example, we could subdivide the edge between *r* and *A* and make *F* the child of the added node. Or we could subdivide edge (*a, b*) and make *F* be the child of the added node. In both cases, the new network will be compatible with the information obtained from the SNPs we have available. Thus, it will be impossible to have a guarantee of returning the correct unrooted topology for *N* from the set of SNPs we have available.

Thus, it is clear that without enough SNPs to cover all the bipartitions of the network, it is not possible to reconstruct the network *N* . This issue affects all methods, not just CUPNS and Gusfield-Estimate-Unrooted, and so is a fundamental limitation for phylogenetic network estimation (and we note that to some extent this is also an issue for phylogenetic tree estimation: if there are no changes of any site on some edge in the true tree, the estimated tree cannot be guaranteed to be accurate). The best that can be done in such situations is to construct, if possible, either one or more phylogenetic networks consistent with the input data, or some kind of consensus network.

### 5.3 Robustness to oracle error

So far, we have assumed that the oracle does not make any errors, so that all sites that are indicated to be SNPs (i.e., homoplasy-free two-state characters) are indeed SNPs, and all sites that are truly SNPs are indicated as such. Thus, the oracle does not make Type I errors (falsely identifying a non-SNP as a SNP) or Type II errors (falsely identifying a SNP as a non-SNP). Here we examine the consequences of oracle error.

We begin with the case where the Oracle never indicates a site is a SNP (homoplasy-free) unless it is in fact a SNP, but may fail to correctly identify some sites as SNPs (i.e., it does not make Type I errors but may make Type II errors). In this case, as long as the probability of failing to say “YES” when a true SNP is presented is less than 1, both CUPNS and Gusfield-Estimate-Unrooted will still be statistically consistent, because in the limit the set *Q_SNP_* (as defined by the oracle) will converge to a set that covers the bipartitions of *N* . On the other hand, if the Oracle has probability 1 of saying “NO” for any SNP defining a particular bipartition in *Bip*(*N_r_*), then there will be incomplete information, and even in the limit the set *Q_SNP_* will not cover the bipartitions of *N* . In that case, as we described in the previous section, both methods will fail.

Now consider the case where the Oracle may indicate that some site is a SNP where it is not. In this case, the set of quartet trees given to CUPNS will include some quartet trees that are not in *Q*(*N_r_*), and this could easily lead to failure. For example, the SN-tree could contain just one large polytomy and no way of resolving that polytomy would lead to the recovery of the network topology. Similarly, the set of bipartitions (characters) given to Gusfield-Construct-Unrooted could lead to the same tree with one large polytomy, and again lead to failure to recover the network topology. Thus, this type of error is much more severe.

In general, these methods are not robust to Type I errors (where a non-SNP is falsely identified as a SNP) and robust to Type II errors (where a SNP is falsely identified as a non-SNP), but only when the probability of Type II errors is not 1 for any given SNP pattern.

### 5.4 Extensions to multi-state characters

So far, we have only examined the question of how to construct a level-1 phylogenetic network from SNPs, which by our definition are two-state homoplasy-free characters. Here we consider the case where the sites that are homoplasy-free can be multi-state (e.g., in a DNA alignment the site might exhibit all four nucleotides). We will show that Gusfield-Construct-Unrooted cannot be applied in this case, but that the CUPNS algorithm is still guaranteed to be statistically consistent.

Recall that Gusfield-Construct-Unrooted creates a character incompatibility graph (i.e., each vertex represents one of the input SNPs, which is treated as a binary character, and edges exist between incompatible characters). Then, the character incompatibility graph is used to identify a set of characters that are compatible with all other characters in the set. Since the characters are binary, pairwise compatibility ensures setwise compatibility (Gusfield, 1991). Therefore, the set of characters that are compatible with all other characters defines a setwise compatible set. For binary characters, a set of compatible characters has a unique compatibility tree (minimally resolved tree that is compatible with all the input characters). Moreover, when the input is a set of SNPs, the compatibility tree is the SN-tree, and is the same tree computed by both Gusfield-Construct-Unrooted and CUPNS.

This approach of computing a character incompatibility graph and then using it to construct a (potentially unresolved) tree that will then be modified to produce a phylogenetic network does not work when the characters are multi-state, because even for three-state characters, pairwise compatibility does not imply setwise compatibility. Furthermore, even if setwise compatibility holds, there can be more than one minimally resolved perfect phylogeny in this case. This also complicates the next step where the polytomies are resolved into cycles, as this too depends on finding a perfect phylogeny using some characters. Therefore, Gusfield-Construct-Unrooted implicitly depends on all the characters being two-state, and cannot be used for multi-state characters.

In contrast, CUPNS is already suitable for multi-state characters! In the first step, it constructs the set *Q_SNP_* of quartet trees based on pairs of states in the SNPs, and so produces quartet trees *uv*|*wx* whenever *c*(*u*) = *c*(*v*) ≠ *c*(*w*) = *c*(*x*). That set of quartet trees is then used to construct the SN-tree and to resolve the polytomies into cycles. All guarantees of correctness, statistical consistency, sample complexity, and runtime follow as for the case where the sites are two-state.

### 5.5 Extensions of the sequence evolution model

In this study, we have assumed that the sites in a multiple sequence alignment evolve down the phylogenetic network *N*, each site picking a single tree in *𝒯_N_* down which to evolve, and then evolving down that tree under a model that allows each edge *e* to have its own substitution probability matrix *p*(*e*; *s, s*′), where *s* and *s*′ are character states. This is a generalization of the General Markov Model (Steel, 1994). Furthermore, we can also allow models of sequence evolution where each site picks a mechanism randomly (e.g., see (Gingerich, 2009)). Heterotachy, where rates of evolution vary with the site and edge (Lopez et al., 2002), represents a kind of heterogeneity that can be problematic for phylogeny estimation; and in its extreme form, the *no-common-mechanism model* (Tuffley and Steel, 1997), can make maximum likelihood estimation statistically inconsistent. Here we ask: do any of the theoretical guarantees we established for CUPNS and Gusfield-Estimate-Unrooted change if we allow the sites to evolve under these more complex models? Interestingly, statistical consistency still holds for these methods under these complex models, since each clade in the network has strictly positive probability of being generated by a SNP character. The argument in each case is surprisingly easy to make, showing clearly that the ability to identify SNP characters (here ensured by availability of a perfect oracle) makes a large difference in the estimation of evolutionary histories, even enabling statistically consistent phylogenetic network estimation.

## 6 Summary and Discussion

The main focus of this study has been on the question of whether unrooted level-1 phylogenetic networks can be constructed correctly from binary (two-state) characters that evolve under the infinite sites assumption, but where we do not know the ancestral state. We establish that given such characters, which we refer to as SNPs, two methods (a previous method presented in Gusfield (2005) and a new method that we develop, called CUPNS) are guaranteed to correctly reconstruct the unrooted topology of the true network as long as it has no cycles of length less than five. To the best of our knowledge, this is the first paper to establish identifiability of unrooted level-1 networks from SNPs, as the proof of correctness for Gusfield’s method has not been previously established. We also establish that these two methods can be used in statistically consistent pipelines for estimating unrooted level-1 phylogenetic network topologies under a stochastic model of DNA sequence evolution we provide, provided that we have access to a reliable Oracle that identifies the SNPs. Furthermore, Gusfield’s method uses *O*(*kn* + *n*^3^) time, where the input alignment has *k* sites and *n* taxa.

We consider the extension to characters that evolve without homoplasy but can be multi-state. We show that for this case, Gusfield’s algorithm is inapplicable (because it depends inherently on two-state characters), while CUPNS maintains its theoretical guarantees when given multi-state characters. Furthermore, assuming access to an Oracle to indicate the homoplasy-free sites, CUPNS can be used in a statistically consistent pipeline for this case.

From a discrete math viewpoint, an interesting contribution of this study is the establishment that the set *Q*(*N*) of all quartet trees in an unrooted level-1 phylogenetic network *N* is a proper superset of the set *Q*(*N_r_*) of quartet trees induced by the rooted trees contained within *N* (i.e.,*𝒯_N_*), provided that *N* has at least one cycle of size five or more. We also establish that a prior method in Gambette et al. (2012) for constructing unrooted level-1 phylogenetic network topologies from *Q*(*N*) fails on *all* level-1 phylogenetic networks *N* with at least one cycle of length at least five, when given *Q*(*N_r_*). This observation represents a cautionary note since SNPs provide information about bipartitions in *Bip*(*N_r_*) and quartet trees in *Q*(*N_r_*), but do not provide evidence for quartet trees in *Q*(*N*) \ *Q*(*N_r_*); methods that aim to estimate phylogenetic networks from data need to depend on what can be inferred from data, rather than on theoretically defined objects that cannot be estimated from the data, such as *Q*(*N*).

Despite the positive results in this study, we also established that CUPNS and Gusfield’s method for constructing unrooted level-1 networks from SNPs without known ancestral state only have strong positive guarantees under idealized conditions: where the input set of SNPs is large enough to cover the bipartitions of the network and the Oracle does not make any mistakes. Thus, both methods are currently only relevant to understanding methods from a theoretical perspective.

One obvious direction for future work is therefore to address the case where the Oracle can make mistakes, and in particular when it can identify some homoplastic sites as being homoplasy-free. Another obvious direction is to design methods that will produce estimates of the phylogenetic network topology even when the input is insufficient to fully define the network. It is possible that heuristics may be developed that are robust to low levels of Oracle errors, homoplastic characters, and incomplete data.

Another question of interest is identifiability of the topology of phylogenetic networks. For example, we are aware of two studies that have examined the identifiability of level-1 phylogenetic networks from sites that evolve down trees within the network, but allowing for homoplasy (Gross et al., 2021; Allman et al., 2022), and another that examined identifiability of level-1 phylogenetic networks from trees that evolve within the network (Xu and Ané, 2023). Identifiability of topologies for tree-child phylogenetic networks (a different class of phylogenetic networks than the level-1 networks we consider) has been examined under a recombination-mutation model of evolution by Francis and Moulton (2018). Statistical consistency is not possible without identifiability, but identifiability in itself does not guarantee that a statistically consistent method is possible—and the development of a statistically consistent method may still be difficult to achieve. Thus, another direction for future work is to see if statistically consistent methods can be developed using the theoretical ideas in these papers.

## Acknowledgments

We thank the reviewers of an early version of this paper (Warnow et al., 2024) for their helpful suggestions.

## Authors’ Contributions

The authors all contributed to conceptualization, formal analysis, writing, review, and editing. Y.T.: theory, original draft preparation. T.W.: theory, project administration, supervision, original and final draft preparation. S.N.E.: theory, original and final draft preparation.

